# Aging the Brain: Multi-Region Methylation Principal Component Based Clock in the Context of Alzheimer’s Disease

**DOI:** 10.1101/2022.02.28.481849

**Authors:** Kyra L. Thrush, David A. Bennett, Christopher Gaiteri, Steve Horvath, Christopher H. van Dyck, Albert T. Higgins-Chen, Morgan E. Levine

## Abstract

Alzheimer’s disease (AD) risk increases exponentially with age and is associated with multiple molecular hallmarks of aging, one of which is epigenetic alterations. Epigenetic age predictors based on 5’ cytosine methylation (DNAm) have previously suggested that biological age acceleration may occur in AD brain tissue. To further investigate brain epigenetic aging in AD, we generated a novel age predictor termed PCBrainAge that was trained solely in cortical samples. This predictor utilizes a combination of principal components analysis and regularized regression, which reduces technical noise and greatly improves test-retest reliability. For further testing, we generated DNAm data from multiple brain regions in a sample from the Religious Orders Study and Rush Memory & Aging Project. PCBrainAge captures meaningful heterogeneity of aging, calculated according to an individual’s age acceleration beyond expectation. Its acceleration demonstrates stronger associations with clinical AD dementia, pathologic AD, and APOE ε4 carrier status compared to extant epigenetic age predictors. It does so across multiple cortical and subcortical regions. Overall, PCBrainAge is useful for investigating heterogeneity in brain aging, as well as epigenetic alterations underlying AD risk and resilience.

## Introduction

Aging is the most significant and consistently demonstrated risk factor for Alzheimer’s disease (AD) [1,2]. As a result, the aging of the U.S. population is expected to coincide with a rise in AD cases, increasing from 5.8 million in 2020, to 13.8 million projected by 2050 [3]. Chronological age, defined as time since birth, is a non-modifiable risk factor. Biological aging however, or the molecular and cellular changes that underlie the process of aging, may be malleable [4–6]. Approaching the challenge of AD prevention and treatment through the lens of biological aging thus provides a major opportunity for improving cognitive health and reducing disease burden.

Because the brain is the central site for AD pathology, understanding the specific aging of this tissue is a priority. As brain tissue ages, misfolded tau and amyloid proteins accumulate due to a loss of proteostasis, one molecular hallmark of aging [7]. While this occurs in older adults with normal cognition or mild cognitive impairment, it is generally more pronounced in subjects with AD dementia [8–10]. Neuritic plaque and neurofibrillary tangles form the basis for the neuropathological diagnosis of AD [11]. Further, more rapid accumulation of tau [12] and β-amyloid [13] aggregates is also linked to inheritance of the APOE ε4 allele, which is itself linked to a number of age-related outcomes, including increased AD dementia risk [14], CVD risk [15,16], and reduced lifespan [17,18]. This suggests that while AD may not be a normal part of aging, it is partially driven by changes that are known to relate to basic aging processes.

Additional hallmarks of biological aging, which include epigenetic alterations [19,20], have also been implicated in the pathology of AD. For instance, 5’ cytosine methylation (DNAm) differences have been shown to track aging and can be quantitatively combined to produce composite aging biomarkers, termed “epigenetic clocks” [21]. We and others have shown that the divergence between observed and predicted ages produced by epigenetic clocks relate to AD pathology. For instance, Horvath pan-tissue [22] and Levine PhenoAge [23] epigenetic age acceleration correlate with abnormally high neuritic plaque, NFT, and β-amyloid loads. While this provides further molecular evidence of a link between AD risk and aging, such clocks are typically developed in peripheral tissues and may not capture the unique aging changes in the brain. A notable exception is the recent DNAmClock_Cortical_[24]. A recent study using Cortical tissue found that the DNAmClockCortical had much stronger associations with AD clinical and neuropathologic traits relative to Horvath, Hannum and PhenoAge clocks based on non-neuronal tissues [25]. Building on previous work, our study incorporates two additional novel features. First, we sought to investigate the extent to which the uneven pathological burden evident by amyloid [26] and tau [27] staging is captured when considering DNAm across multiple paired brain regions, rather than in singular areas of the cortex or hippocampus. Second, we have previously shown that CpG clocks often suffer from significant technical noise, hindering their applications. Therefore, we sought to use our recently developed approach to improve signal-to-noise ratios in methylation data, leading to improved reliability and construct validity in our novel epigenetic clock [28].

Overall, we hypothesized that a brain age methylation-based predictor could be developed with meaningful disease associations and broad multi-brain-region utility. To test this, we used DNAm capture to generate a PC-based epigenetic predictor of brain aging which we show to: (1) strongly reflect AD neuropathology and cognitive decline, and (2) track age across multiple brain regions. This resulting measure, PCBrainAge, is applicable for use in existing brain and tissue banks, and many publicly available postmortem datasets for the study of AD.

## Results

### Model Design & Testing

To generate a predictor of aging in the brain, we selected a publicly accessible dataset deposited into the Gene Expression Omnibus [29] (GSE74193) [30]. In brief, this dataset contains methylation data from dorsolateral prefrontal cortex (DLPFC) of 399 individuals aged 20+ (see Methods for further details). This dataset includes patients diagnosed with schizophrenia (n = 187; 47%). However, this neuropsychiatric disease has not been shown to be robustly associated with epigenetic signatures of chronological age in either blood or brain, despite acceleration in clocks predicting mortality in blood [31,32]. Our model’s outcome variable is chronological age, so inclusion of schizophrenia samples is reasonable and potentially advantageous: Inclusion of schizophrenia samples reduces the likelihood that general brain pathology will exert a large impact on the model’s predictions, as the model is forced to predict chronological age despite schizophrenia status. Nevertheless, as a sensitivity analysis, we also trained a model using only control individuals, which did not improve results (Figure S1). Thus, we included all high-quality samples for training, regardless of schizophrenia status.

The training method to generate our predictor is built upon our recently published PC Clock method [28]. In brief, singular vector decomposition (SVD), an extension of principal components analysis suitable for wide format data (i.e. where features outnumber samples), was performed on this training methylation dataset. This analysis was limited to CpG sites that are overlapped between the training, test, and validation datasets collected on 450K or EPIC arrays (see Methods for more detail). This produced 399 left singular vectors, which for general purposes, are referred to here as principal components (PCs) of 5’-cystosine methylation in postmortem dorsolateral prefrontal cortex (DLPFC). The PC scores, representing an individual’s projection values onto the principal component vectors, were used as the set of variables from which age was predicted via elastic net penalized regression.

To predict training sample age, three models were generated, differing in the sex representation of subjects. This choice was based upon known sex specific differences in aging [33], and evidence of sex-specific differences in AD risk and AD neurobiology [34], and. All models used elastic net penalized regression in the appropriate individuals to find the optimal weighted linear average of PCs to predict chronological age; the first utilized both sexes (n = 399, Figure 1A,D); the second was fit to only males (n = 262, Figure 1B,E); the third was fit to only females (n = 137, Figure 1C, F). Regardless of the sample used for training, we found that each model attained similar correlation between predicted and chronological age in both males and females. Male- and female-specific age correlations for each model can be found in Table 1.

**Figure 1:**
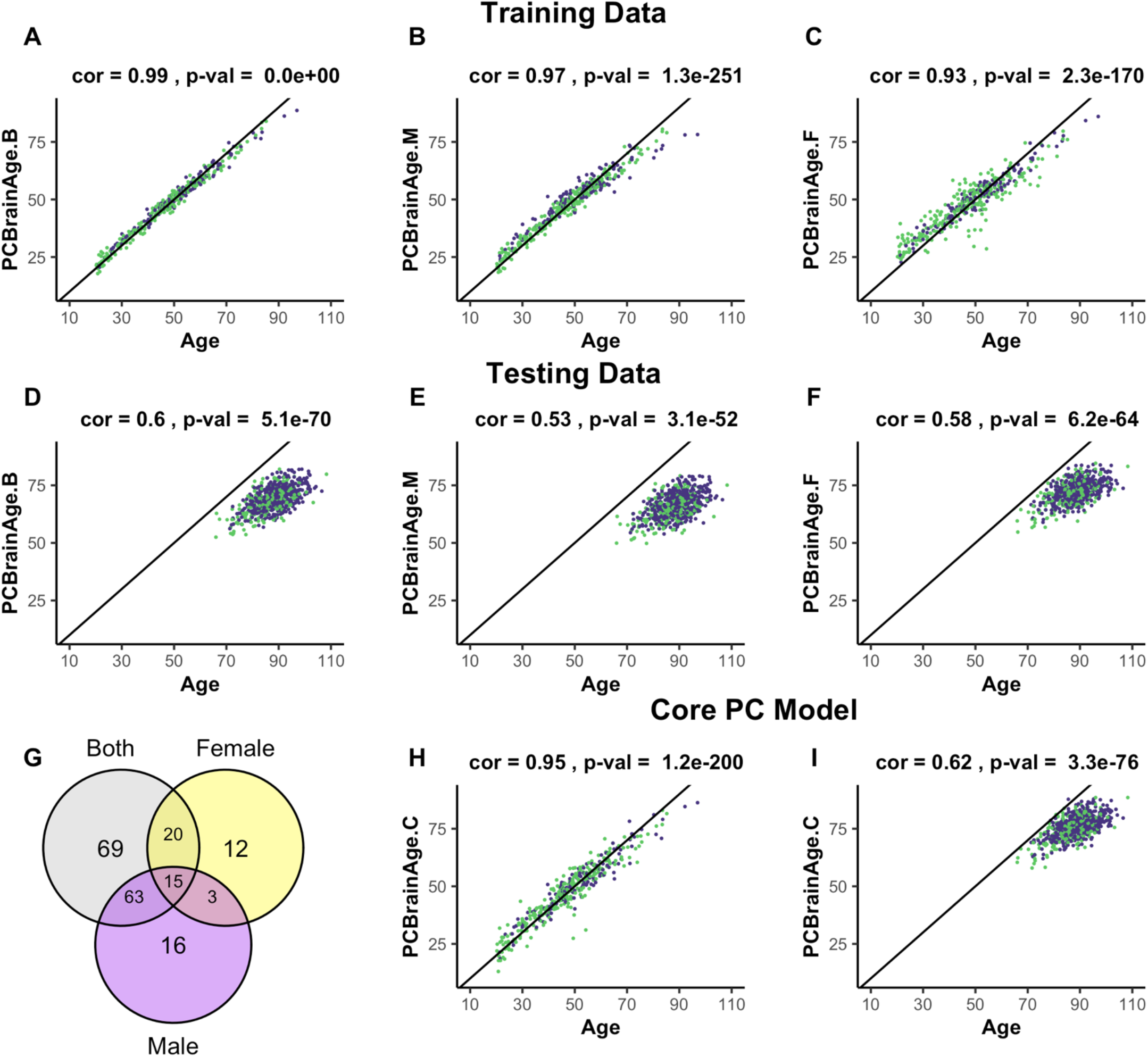
Training and testing of multiple iterations of PCBrainAge. Using the dataset from **GSE74193**, elastic net was used to predict age using principal component loadings in both sexes (**A**), only males (**B**), or only females (**C**). Here, we show the resultant predictions for each model in both females (purple) and males (green) regardless of training sex. Each model so trained is then predicted in all individuals from **syn5850422** (**D-F**), regardless of sex or AD status. Each model selected a number of principal components to use for prediction, and we compared the selection of each model using a Venn diagram (**G**). Subsequent training of an elastic net model using only the 15 core principal components in both sexes is visualized (**H**) and compared to performance in the test dataset (**I**).

**Table 1.** Pearson’s Correlation of Age by Sex with Model Age Predictions

| Model |  | Training Data | Core | Test Data | Core |
| --- | --- | --- | --- | --- | --- |
| Both Sexes | M | 0.99 | 0.94 | 0.61 | 0.65 |
|  | F | 0.99 | 0.95 | 0.56 | 0.58 |
| Male | M | 0.98 |  | 0.55 |  |
|  | F | 0.95 |  | 0.49 |  |
| Female | M | 0.90 |  | 0.61 |  |
|  | F | 0.97 |  | 0.54 |  |

A total of 195 of 399 PCs were selected for use in one or more models, with the female model (PCBrainAge.F) selecting the fewest variables. However, a completely overlapped set of 15 PCs (referred to as the core PCs) were selected for all three models, representing an important centralized signal of aging (Figure 1G). The creation of three degenerate models in this manner allowed us to isolate a robust brain aging signal. We investigated whether these core PCs were significantly more important than their non-core counterparts in each model. To do so, we sequestered the 15 core PCs in the training data, as well as 3 different sets of non-core PCs corresponding to each original model. Using the same elastic net regression procedure as the original models, we regressed the core and noncore PC scores to age in the appropriate training subcohorts. The 15 core principal components were sufficient to predict age in the training data. However, the non-core models were unable to successfully do so (Figure S2). Therefore, we generated a final model PCBrainAge.C. This was trained in both males and females and includes only the 15 core PCs (Figure 1H). As the core PC version of PCBrainAge is the overall superior model—both requiring limited information and performing best—only this model is used hereafter and will be simply referred to as PCBrainAge for clarity.

To validate models of aging generated from training data, an independent methylation dataset of 718 DLPFC samples was obtained through Synapse (syn5850422) [35]. All datasets used in the current work are characterized in Table S1. Estimation of the individuals’ PC loadings was performed by projection onto the right singular vectors of the training dataset, thereby generating the 399 training PC vectors based upon the original eigenvalue estimations from the training data [28]. In terms of age predictions, PCBrainAge.C performs at least as well as all original sex-stratified and full models in the test dataset (Figure 1I), despite using fewer principal components.

Principal components are complex, composite variables, making them challenging to interpret. To investigate the information captured in the 15 core PCs, we correlated each PC to annotated features of the training dataset. This demonstrated that the largest source of variation in the data, as captured by PC1, is cell composition (Figure 2A). PC5 is most strongly correlated with age (|r| = 0.68), with all PCs having a range of absolute biweight midcorrelation of 0.04-0.68. As the overall model has correlation with age of 0.95, the signal for chronological age is clearly distributed across PCs. PC8 and PC15 are related to biological sex. These observations were confirmed when the PCs were projected onto the test data (Figure 2B). While neuron proportion demonstrates correlations with more PCs in the test data compared to training data, there are several explanations for this observation. First, it is consistent with an expected, subtle loss of the imposed orthogonality when principal components are projected into new datasets. Second, our test dataset is comprised of only older adults, many with AD, who are expected to have age-associated neuron loss. This may enhance correlations between cell composition and methylation PCs that were not otherwise apparent in training data capturing the entire lifespan. Third, the proportion of neurons in test data, unlike training data, is estimated using the methylation itself. Therefore, this estimated cell proportion may be doubly affected by disease states or other signals being captured in the data. Finally, in the test dataset, where some individuals have AD and dementia, we find that no single PC is highly and/or consistently correlated to AD status (Table S2).

**Figure 2:**
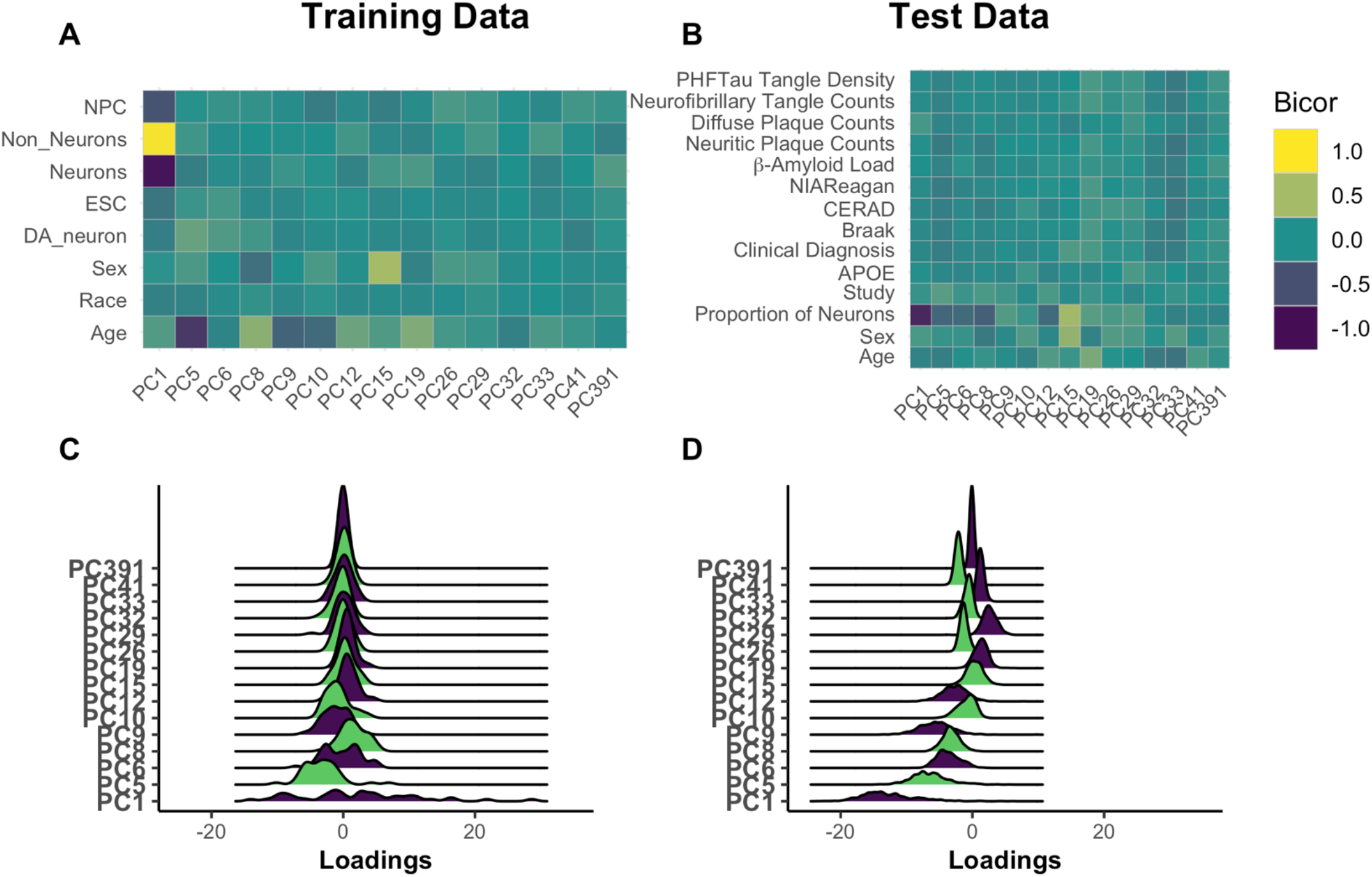
Understanding Core Principal Component Composition. Principal component loadings for individuals in the training dataset were correlated using biweight midcorrelation (bicor) to selected author-provided phenotypic annotations (**A**). The same procedure was applied to the projected principal component loadings for all individuals in the test dataset, including those with and without Alzheimer’s disease (**B**). To ensure that future correlations between age prediction and disease are not the result of unrealistic distortions in PC loadings following the prediction process, we used ridgeplots to visualize the distribution of loadings in each PC in age 65+ training individuals (**C**) and the test data (**D**). [Abbreviations: NPCs—neural progenitor cells; Cort.—cortical; ESCs— embryonic stem cells; DA—dopaminergic; np—neuritic plaque]

We checked for relative agreement between training and testing data composition prior to applying PCBrainAge to test data. We previously showed that it is possible to use PCs from one dataset to generate reliable and useful PCs when projecting to another [28]. However, there can be differences in the distributions of PC scores: Therefore, we analyzed the distribution of PC scores from the core and found that while there are shifts in the mean, the general shape remains intact (Figure 2C,D). This can contribute to differences in the intercept in new datasets, a known behavior in common CpG clocks. However, this is easily corrected by using age acceleration, which does not consider intercept, for further analyses.

### PCBrainAge Correlates with Alzheimer’s Pathology in DLPFC

We calculated brain age acceleration in the ROSMAP DLPFC test dataset (n = 700) by generating linear models to regress PCBrainAge on samples’ true ages at death and proportion of neurons (Table S3). Proportion of neurons was explicitly included to obtain residuals, as we hypothesized that cell proportion changes appear to be the dominant signal in data (Figure 2A, B). Ultimately, we are interested in whether PCBrainAge is predictive of AD beyond the well-characterized impacts of changes to neuron abundance. This is also consistent with previous reports that correcting for cell type heterogeneity improves mortality and biological age prediction [36,37]. Age acceleration was correlated with pathological and phenotypic traits known to indicate or affect the course of AD. To ensure that such correlations were not impacted by a nonlinear relationship of age and PCBrainAge, we verified a uniform distribution of residuals in both sexes (Figure 3A). Slight nonlinearity at the extremes of the distribution appeared to be the result of reduced sample density rather than true nonlinearity.

**Figure 3:**
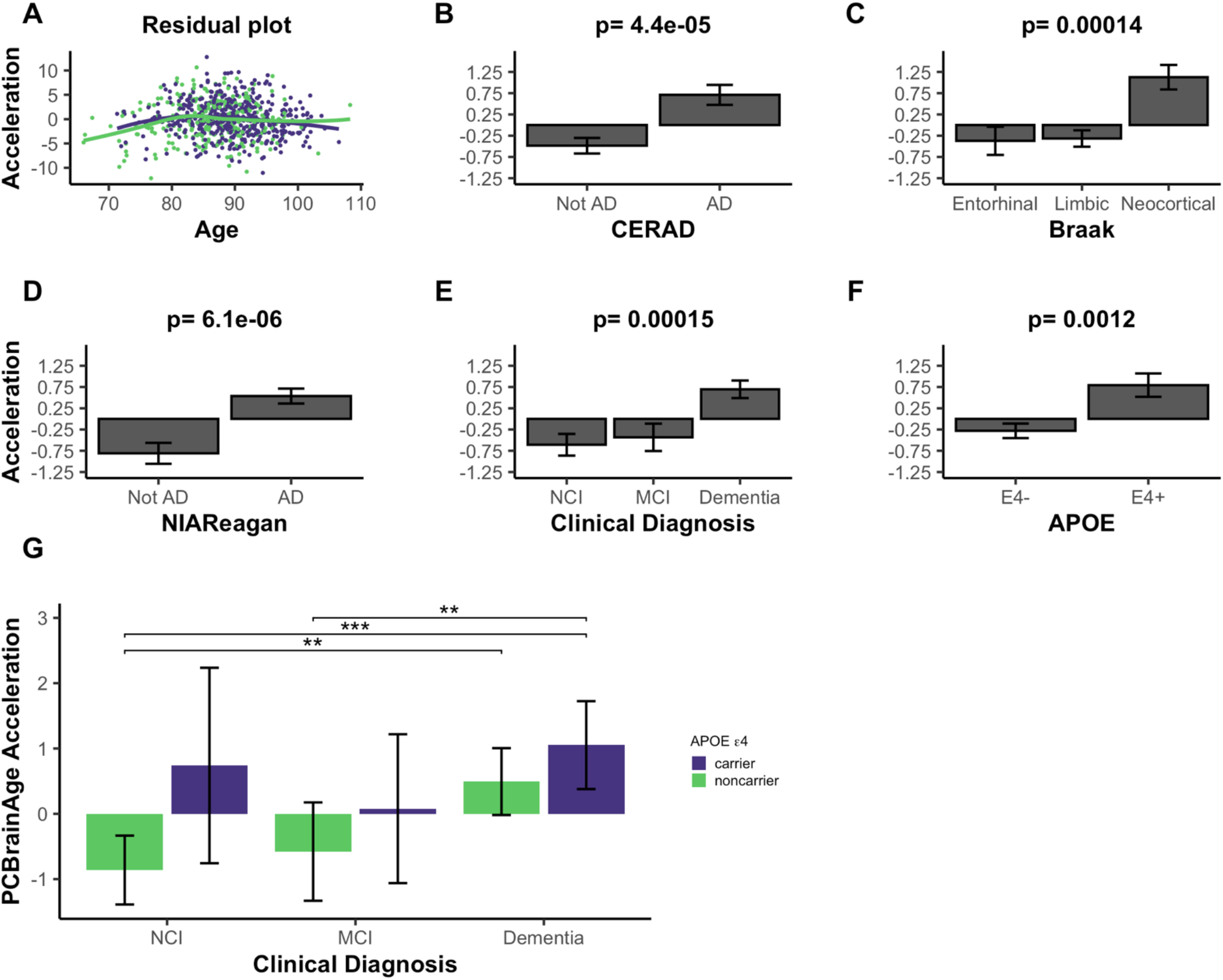
PCBrainAge Acceleration is Associated with Indications of AD. (**A**) PCBrainAge residuals following multiple correction were verified to remain orthogonal to age using a scatterplot with LOESS curves for males (green) and females (purple). PCBrainAge Acceleration was subsequently analyzed in the context of CERAD scores (**B**), Braak stages (**C**), NIA Reagan scores (**D**), the antemortem clinical diagnoses (**E**), and the APOE ε4 carrier status (**F**) of each individual. P-values are the result of performing Kruskal-Wallis tests of nonparametric means amongst the categorical groups. Error bars for 3B-3F depict 1 standard error. (**G**) Acceleration was further broken down into cognitive groups by APOE ε4 carrier status for improved clarity. Error bars depict the 95% confidence interval. Significance levels based on BH adjusted p values are: * < 0.05, ** < 0.01; *** < 0.001.

PCBrainAge accelerations were tested for association with AD clinical and pathologic diagnosis, and APOE ε4 status. Postmortem binarized AD diagnosis according to neuritic plaque derived CERAD scores is significantly associated with accelerated brain aging (Figure 3B), as well as neurofibrillary tangle (NFT) derived Braak Staging (Figure 3C). Notably, PCBrainAge is significantly accelerated when AD is in the neocortical, final stages versus all prior stages, but has no discriminatory power between the entorhinal and limbic phases. This likely reflects that PCBrainAge’s training and testing are occurring in DLPFC, a neocortical region. PCBrainAge acceleration is also positively associated with post-mortem neuropathologic AD diagnosis by combined neuritic plaque (np) and NFT to derive NIA Reagan score (Figure 3D), as well as ante-mortem clinical diagnosis of AD dementia (Figure 3E). Those with the clinical diagnosis of mild cognitive impairment (MCI) are indistinguishable from non-cognitively impaired individuals. Thus, individuals with AD show greater PCBrainAge acceleration than their counterparts. The APOE ε4 allele has been reproducibly associated with AD risk, and earlier onset of the disease [38]. Positive APOE ε4 status (i.e. carrying 1 or 2 APOE ε4 alleles) was significantly associated with PCBrainAge acceleration (Figure 3F). In fact, PCBrainAge is accelerated across APOE ε4 carriers such that cognitively normal and AD confirmed individuals are indistinguishable. In contrast, among non-carriers those with AD show significant acceleration over premortem cognitively normal individuals (Figure 3G).

During the course of the current research, another methylation-based epigenetic clock was reported, termed DNAmClock_Cortical_ [24]. As expected, DNAmClock_Cortical_ is correlated with PCBrainAge for both predicted age and age acceleration (r=0.79 and r=0.56, respectively) (Figure 4A, B). The training samples of DNAmClock_Cortical_ (n = 1,047) included the samples used to train PCBrainAge (n = 399) and was intended to accurately estimate chronological age of samples at all ages (training and test samples aged 1-108). Indeed, DNAmClock_Cortical_ does show better correlations with chronological age compared to PCBrainAge. However, our work with PCBrainAge intends to not only predict age, but to capture relevant biological heterogeneity of aging in the brain—especially that associated with AD. DNAmClock_Cortical_ acceleration shows less significant associations with AD clinical and pathologic phenotypes, and APOE ε4 carrier status in comparison with PCBrainAge (Figure 4C-G compared to Figure 3). This then suggests that DNAmClock_Cortical_, in optimizing prediction of chronological age, may miss relevant heterogeneity and aging signals associated with AD. A 1-SD change in DNAmClock_Cortical_ acceleration does correspond to odds of pathologic AD, but this is mostly limited to amyloid and neuritic plaques. However, a 1-SD difference in PCBrainAge reflects greater differences with AD pathology, and is more balanced across various postmortem metrics of AD pathology (Figure 4H). Further, it was found that increasing standard deviations of PCBrainAge acceleration show monotonic increases in the normalized probability of dementia, unlike the stochasticity observed in DNAmClock_Cortical_ (Figure 4I). This is likely reflective of the hypothesized reduction in noisy CpGs and improved resolution expected to arise from using our PC Clocks method.

**Figure 4:**
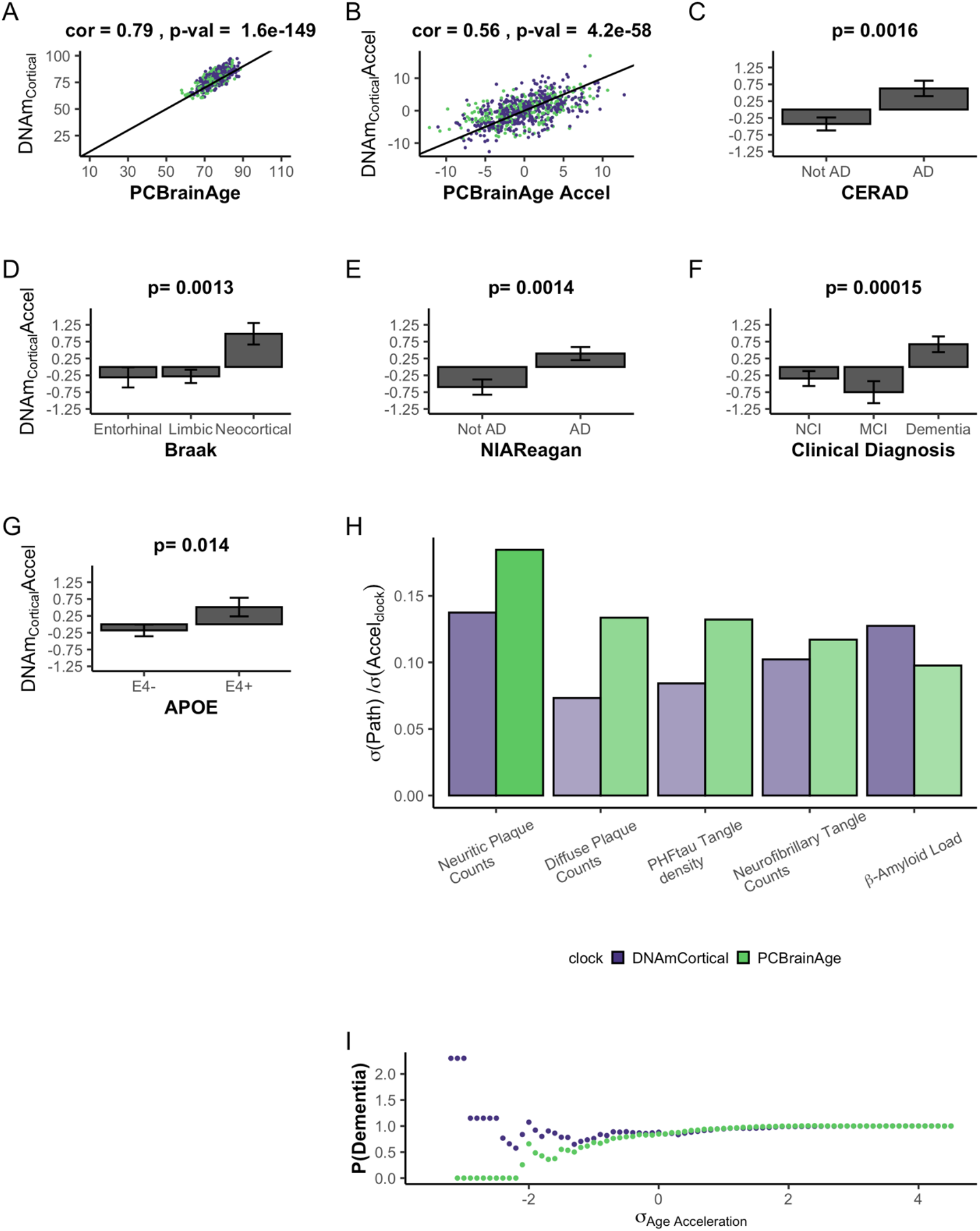
DNAmClock_Cortical_ Prediction in Test Data Comparable to PCBrainAge Predictions. DNAmClock_Cortical_ was estimated in our test dataset, which is independent from it’s original training. We find that DNAmClock_Cortical_ has moderate correlation with age at death (**A**), and agreement with PCBrainAge accelerations for the same individuals (**B**). While DNAmClock_Cortical_ does exhibit clear acceleration in (advanced) AD patients (**C-E**), demented patients (**F**), and APOEε4 carriers (**G**), the p-values of the separation between groups are slightly attenuated versus those of PCBrainAge (see Figure 3). The standard deviation of various AD pathological characteristics per clock standard deviation are compared for DNAmCortical (pink) and PCBrainAge (blue) (**H**). Given individuals less than or equal to a standard deviation of age acceleration for each clock, the probability of patients being diagnosed with dementia normalized to the total cohort probability is shown for each clock (**I**).

The association of PCBrainAge acceleration and AD pathology suggests a discriminatory role for PCBrainAge beyond age prediction itself. Increased prediction accuracy of chronological age may reduce the association with AD. PCBrainAge provides meaningful, nonrandom information about both age and the disease status of the brain.

### Alzheimer’s Pathology Correlates with PCBrainAge Across Multiple Brain Regions

Aging may have distinct effects on different brain regions with respect to atrophy, dendritic morphology, synaptic plasticity, and vasculature [39]. This may be reflected in epigenetic clocks, which indicate measurable differences in brain aging rates between regions [40]. The typical progression of AD involves the reproducible, staged invasion of neurofibrillary tangles [27] and amyloid-β aggregations [26] through the brain. Though AD progression can be variable, some regions show amyloid or tau pathology earlier than others [41]. It is unknown whether regional differences in epigenetic age might help explain the differential impact of AD pathology amongst brain regions.

We used PCBrainAge to measure the aging trends across multiple brain regions and evaluate regionspecific associations with AD. Using 333 individuals’ samples from an APOE ε4 carrier enriched subcohort of ROSMAP (Table S1), we generated novel DNAm data from 3 distinct brain regions for each individual: Prefrontal cortex (PFC), Striatum (ST), and Cerebellum (CBM). This incorporated 212 overlapped individuals for which DNAm data for DLPFC (a distinct region and tissue slice) was available in the original test dataset used here. As done for the original test dataset, principal components were projected into this data, followed by PCBrainAge prediction for each independent region and sample. To account for repeated measurements and to improve modeling of epigenetic age acceleration, we employed a linear mixed effects (LME) model to utilize data across regions in tandem. The model is described in equation 1. The three brain regions tested here are expected to diverge in their epigenetic age prediction based upon data in prior clocks [42–44]. Therefore, we allowed a random effect to the model intercept with age according to brain region. Comparison of this model and simple regression is shown in Table S4.

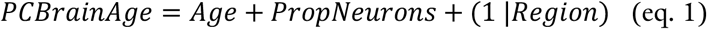

We then related age acceleration in each region to AD neuropathology, clinical and pathologic diagnoses, and APOE ε4 carrier status. To account for multiple comparisons, we used a p-value adjustment according to a Benjamini Hochberg procedure from a Kruskal Wallis test of nonparametric mean differences (Table 2). Age acceleration in prefrontal cortex and striatum were both associated with AD neuropathology and premortem clinical diagnosis. Age acceleration in the striatum was also higher in accordance with APOE ε4 carrier status (Figure 5). It is important to note that the dataset used here is a subset of the test dataset used in Figure 3 (See Figure S5). The associations here are weaker compared to Figure 3 likely due to the reduced power, as this dataset contains fewer samples. This effect is magnified when comparing multiple brain region samples for each individual which increases complexity of comparisons without an increase in the number of individuals.

**Table 2.** Multi-Region PCBrainAge Acceleration’s Correlates to AD

|  | All | PFC | ST | CBM |
| --- | --- | --- | --- | --- |
| **CERAD** | 1.1E-04* | 6.3E-03* | 6.3E-03* | 0.13 |
| **Braak** | 0.087 | 0.25 | 0.067 | 0.15 |
| **NIA-Reagan** | 4.5E-04* | 6.3E-03* | 6.3E-03* | 0.36 |
| **Clinical Diagnosis** | 6.3E-03* | 6.3E-03* | 4.7E-03* | 0.13 |
| **APOE** | 0.24 | 0.52 | 0.076 | 0.68 |

\* indicates a p-value < 0.05 of Kruskal Wallis nonparametric tests of means across criterion groups, followed by adjustment according to the Benjamini Hochberg procedure when pooling all p-values in the table

The cerebellum has long been characterized as relatively spared in AD, though this has been challenged recently [43]. Interestingly, the cerebellum ages slowly according to the multiple epigenetic clocks, and existing epigenetic clocks do not show correlations between cerebellum age acceleration and AD neuropathology [42] (Figure S4). These other clocks were not trained in brain tissue, so it remains plausible that brain—or even cerebellum specific—epigenetic aging signatures are correlated with AD. However, we found that PCBrainAge acceleration in cerebellum is not significantly correlated with AD diagnosis, neuropathology, or APOE ε4 carrier status (Figure 5). Thus, PCBrainAge validates prior reports that the cerebellum’s methylation age diverges from that of other brain regions reflecting its distinctive biology in AD.

Taken together, PCBrainAge demonstrates associations with AD neuropathology, diagnosis, and APOE ε4 carrier status in three regions affected by AD (DLPFC, PFC, and striatum) but not in a region that may be relatively spared in AD (cerebellum). Furthermore, PCBrainAge is applicable in multiple brain regions despite being trained specifically in DLPFC.

## Discussion

While epigenetic clocks trained in blood, or multiple tissues, can reflect age in brain tissue [45], biomarkers trained specifically for the brain may more accurately capture its aging trends. Clocks trained in peripheral tissue can reflect postmortem AD pathology when applied to brain DNAm data [46]. However, with the exception of the PhenoAge clock [23], the acceleration captured does not typically demonstrate significant association with AD dementia status, despite clear correlations with neuropathologically mediated cognitive decline [46]. This may reflect the intermediate complexity between molecular pathological change and higher order cognitive changes [reviewed in 45]. However, blood or pan-tissue trained clocks may not adequately capture brain aging, due to the brain’s unique methylation profile [48–50], extreme diversity of specialized neuronal [51] and glial [52] cell types, and distinct developmental patterns [53]. Thus, a methylation-based predictor of age in the brain is useful for studying age-related patterns of change in neurodegenerative disease at its source.

Prior work has been done to develop methylation-based predictors of age in the brain in humans [24] and mice [54]. A biomarker of human brain aging, DNAmClock_Cortical_, addresses the systematic underestimation of age in older adults when predicting brain age by existing clocks. While DNAmClock_Cortical_ can achieve near-perfect age correlation in brain tissue, this was not the goal of the present model. The lower correlation of PCBrainAge in the test datasets, as depicted in Figure 3, carries important biological signal. While the present clock does not achieve the degree of correlation of clock age and sample age at death found by DNAmClock_Cortical_, PCBrainAge’s utility lies in the robust link between an individual’s PCBrainAge residual (age acceleration), and pathological characteristics of AD. Beyond this, generation of PCBrainAge employed a novel methodology that allowed few samples for adequate training and was shown to reduce technical noise, thereby improving confidence in biological interpretation of the reported age residuals. Reduction of technical noise in this manner has also been hypothesized to reduce the sample size needed to train a robust epigenetic clock model [28], addressing the marked scarcity of brain tissue in comparison to blood DNAm. Furthermore, PCBrainAge demonstrates applicability across brain regions.

While a connection between DNA methylation and AD neuropathology has been a previously discussed possibility [47,55], the current work also demonstrates a connection between patterns of DNA methylation change and higher order changes like those to cognition, and significant genetic differences like APOE ε4 status. The acceleration predicted by PCBrainAge is correlated with clinical AD dementia, and pathologic AD, outperforms sex specific and pooled sex models in both males and females, and can be used across multiple cortical and noncortical brain regions. PCBrainAge is also significantly associated with APOE ε4 status (see Figure 3F), which has not been previously shown with existing blood-based clocks. With APOE ε4 carriers exhibiting PCBrainAge acceleration over their non-carrier counterparts, PCBrainAge is consistent with observations that this genotype significantly increases risk in an age-related manner [56]. PCBrainAge can also detect the interaction between APOE status and cognitive diagnoses, given that APOE carriers show acceleration regardless of diagnosis, while noncarriers with AD show distinct acceleration versus those who appear cognitively normal. One limitation, however, is that our dataset shows an enrichment of APOE ε4 carriers with MCI and dementia over cognitively normal counterparts. Regardless, this may reflect APOE ε4 carriers’ increased neuropathological burden [57,58], while suggesting that APOE carriers may not be aggressively predisposed to higher order cognitive changes.

PCBrainAge can predict age across multiple brain regions while also capturing heterogeneity relevant to AD in that region. The degree of correlation recapitulates previously described differences in the rate of aging of brain samples [40,42]. DLPFC, PFC, and ST are routinely impacted by AD pathology, unlike cerebellum [59–61]. We found that PCBrainAge acceleration is associated with AD pathology and dementia status in these regions. This signal is slightly more robust in ST, where age correlation is stronger and separation between pathological groups is more distinct. It has been well characterized that tau and amyloid impact brain regions at varying times and to varying degrees. Further investigation is necessary as to whether a model of epigenetic brain aging reflects a relationship between pathological temporality and epigenetic alterations.

In the cerebellum, PCBrainAge recapitulates aging deceleration reported in previous studies [42]. Here we also show that age acceleration of cerebellum lacks correlation with Alzheimer’s pathology and disease status. The slower predicted rate of aging in CBM conforms to expectations that CBM aging and its relationship to AD are drastically different from other brain regions. Without knowing the causal direction for the link between age related 5mC changes and AD pathology, the mechanisms for this relationship remain unclear. There is some evidence that amyloid beta can reduce methyltransferase activity resulting in global hypomethylation, and cerebellum is relatively spared of amyloid pathology until very late in the disease [62]. Future studies should investigate these mechanisms.

PCBrainAge is a promising predictor of regional brain aging, with demonstrated recapitulation of known aging trends in multiple brain regions. However, beyond tracking the relative aging of various brain regions, it can assess meaningful age-acceleration, or pathological aging. This pathological age acceleration is further correlated to AD neuropathology, clinical AD diagnosis, and APOE ε4 carrier status. PCBrainAge may aid in future investigations linking heterogeneity in the aging process to AD risk and individual resilience.

## Methods

### Selection of Available DNA methylation Data

DNA methylation data was acquired from multiple sources (Table S1). The training data was accessed from the Gene Expression Omnibus (GSE74193) [30] as the age range was much wider than in the Alzheimer’s Cohort studied: This has the important effect of increasing the ratio of the range of the variable of interest (age) versus the signal (DNAm) noise due to technical error, biological heterogeneity, and the effect of diseases. All sample methylation β values were generated from the dorsolateral prefrontal cortex (DLPFC) using the Infinium HumanMethylation450K Beadchip (Illumina, San Diego CA, USA) and were used as collected, normalized, and reported by the original authors [30]. Samples under the age of 20 from the original GEO dataset were excluded as it has been shown that development typically has a different aging regime when considering epigenetic clocks [22,63].

For assessment of PCBrainAge in the context of neurodegeneration and AD, we used the previously collected synapse dataset (syn5850422) [35] which generated Illumina 450K methylation data in postmortem DLPFC of participants in the Religious Orders Study and the Rush Memory & Aging Project (ROSMAP) [64]. Methylation β values were used as originally collected, normalized, and reported by the original authors [65]. Samples were excluded if their clinical diagnosis value was a non-AD primary cause of dementia [66]. Use of these samples would introduce significant uncertainty beyond the scope of the current work. Clinical diagnoses were dementia, mild cognitive impairment, and no cognitive impairment, and Alzheimer’s dementia proximate to death (n = 700) [67]. Neuropathologic data included CERAD, Braak, and pathologic AD by NIA-Reagan [68]. AD neuropathology was previously generated for this dataset: Neuritic and diffuse plaques, and neurofibrillary tangles were estimated using count data from silver stain; PHFtau tangle density and β-Amyloid load were each estimated using molecularly specific immunohistochemistry [69].

### Generation of Multi-Region Brain Methylation Data

Novel collection of multi-region brain 5 ‘-cytosine DNA methylation data was performed for the current work. This data was collected from frozen brain tissue samples obtained from Rush University’s Religious Orders Study and Rush Memory and Aging Project (ROSMAP) [64]. Frozen tissue was isolated in 349 individuals across three brain regions: Brodmann Areas 10 (prefrontal cortex), 22 (striatum), and cerebellum.

Bulk genomic DNA was extracted from each tissue sample using the Chemagic DNA Tissue100 H24 prefilling VD1208504.che protocol (Perkin Elmer Ref# CMG-1207). In brief, tissue was lysed overnight at 56 degrees in 1mL Chemagic Lysis buffer and 50uL Proteinase K. Samples were treated with 80uL of RNASE A @ 4mg/uL (AmericanBio Ref# AB12023-00100) for 10 minutes at 56C. Lysis was then transferred to a deep well plate and the extraction performed via the Perkin Elmer Chemagic 360 extraction instrument. Samples were centrifuged at 13000RPM for 1 minute, placed on a magnet and transferred to final 1.5mL Eppendorf tubes. 25-50 mg of extracted DNA per sample was then used according to the manufacturer’s protocol on the Illumina Methylation EPIC array at the Yale Center for Genome Analysis (YCGA) with sample randomization on each array to mitigate batch effects.

The raw .idat files of bisulfite-converted single-CpG resolution of methylation were processed to obtain β values through ratios of probe intensities, according to standard methods. Using the R ‘minfi’ package [70], Noob normalization was performed on β values. For more information, the method used herein was derived from a prior publication [71]. The raw (syn23633756) and normalized beta value (syn23633757) data have been deposited to Synapse (Sage Bionetworks, Seattle, WA).

The sample phenotype data was provided as previously generated by the Rush University Alzheimer’s Disease Center, and in accordance with prior publications: Individuals’ clinical diagnoses [67] and neuropathologic data [68] were annotated as in the test dataset.

### CpG Selection

5’-cytosine DNA methylation was collected on two different arrays across datasets: The 450K and EPIC arrays. Therefore, DNAm was limited to only the intersection of sites between the two arrays. Further, CpGs located on sex chromosomes, as indicated in the Illumina 450K array manifest were excluded. This resulted in retention of 357,852 CpGs.

### Model Training

Singular vector decomposition was implemented using the *prcomp* function in the R stats package (v3.6.1). Detailed methods for training and principal component projection can be found in the methods of Higgins-Chen et al. [28]. In brief, centered principal component scores for individuals, understood as an individual’s score based upon their CpG β values undergoing a rotation according to the rotation matrix, are used as inputs for an elastic net regression to predict age at death. That is, rather than using the original beta values for a given individual’s CpGs, the left singular vectors (PCs) are used instead (excluding only the last PC). Elastic net regression was performed using the *glmnet* package in R, according to a mixing parameter (**α**) of 0.5, utilizing equal parts LASSO and ridge regression with 10-fold cross validation. All fit models were estimated using penalization (**λ**) corresponding to predicted minimal error. Training of the age predictor was performed thus in an unsupervised manner, and in a dataset without clinical AD individuals. The final PCBrainAge model, which constitutes the “core” model entailed retraining an elastic net model such that PCs were zeroed out if not one of the 15 core principal components which had some nonzero weight in the original three models.

### Estimation of Neuron Proportion

Along with methylation data, publicly available dataset annotations provided sorted cell proportion estimates, or estimates of neuron and glia proportion calculated using the methylation-based Cell Epigenotype Specific Model (CETS) for R package [72]. In the novel, multi-region dataset, the CETS package was used to estimate the proportion of neurons in each brain region sample.

### Statistical Measures

All reported scatterplot and predictor of age at death correlations (and corresponding p-values) are the result of correlation tests between means according to the Pearson’s product moment coefficient, presuming standard normal distributions. This is implemented using the R function *cor.test* from the stats package. Correlations with annotations of phenotype, as in Figure 2A-B are the result of implementing a biweight midcorrelation, a median-based comparison test that improves sensitivity to outliers. This was implemented using the *bicor* function of the WGCNA package in R.

The p-value reported for all barplots are the result of a Kruskal-Wallis Rank Sum Test, which is a nonparametric test of means. This did not require assumptions of normality, and was applied using the *kruskal.test* function in the R stats package. Error bars on all barplots represent a standard error of the mean, unless explicitly noted otherwise.

In all tables offering many independent p-value comparisons (Table 2,S2), adjustment of p-values were necessary. All adjusted p-values were reported following implementation of the Benjamini Hochberg procedure, and values p <0.05 were considered significant.

Linear mixed effects (LME) models were used in the context of the multiregion data. This required implementation of the *lmer* function of the lme4 package in R [73]. LMEs were optimized according to a Nelder Mead optimizer, and were visualized using the *sjPlot* package. Generation of LMEs were done in a hypothesis driven manner as described in the results, and were compared to the marginal R^2^ of less complex models. Visualization of the model table was performed using sjPlot [74].

## Author Contributions

KLT, AHC and MEL jointly conceived of the current project and contributed to the writing of the manuscript. KLT carried out the analysis with direct feedback and additional suggested analyses from AHC and MEL. CVD and AHC provided clinical perspectives for the current work. SH and MEL contributed background and experience to the conceptualization of the project. DAB and CG provided specimens for data collection, generated clinical, neuropathologic and DNAm data, and critically reviewed the manuscript.

## Acknowledgements

We are grateful to participants in the Religious Orders Study & the Rush Memory and Aging Project, without whom the data supporting the current work would not have been possible. We would also like to thank the NIA’s Resilience-AD consortium for their support and feedback on this work.

