## Supplemental Data 1 for "Aging the Brain: Multi-Region Methylation Principal Component Based Clock in the Context of Alzheimer’s Disease"

#### Supplemental Tables

**Table S1.** Dataset Access, Annotation and Composition

| | Training | Testing | Validation |
|---|---|---|---|
| **Accession** | GSE74193 | syn5850422 | syn23633757 |
| **n** | 399 | 700 | 333 |
| **Array** | 450k | 450k | EPIC |
| **Brain Region** | DLPFC | DLPFC | Matched PFC, ST, CBM |
| **Ages** | 20-97 | 66-108 | 66-108 |
| **Has AD?** | No | Yes | Yes |
| **White** | 0.48 | 1 | 0.97 |
| **Female** | 0.34 | 0.64 | 0.65 |
| **APOE ε4** | N/A | 0.26 | 0.71 |

**Table S2. Biweight Midcorrelation of Core PCs and AD characteristics**

| | Age | p | Cerad | p | Braak | p | NIA-Reagan | p | Cogdx | p | APOE ε4 | p |
|---|---|---|---|---|---|---|---|---|---|---|---|---|
| PC1 | **-0.10** | * | -0.07 | | -0.05 | | -0.04 | | -0.07 | | -0.01 | |
| **PC5** | **-0.14** | *** | **-0.13** | ** | **-0.13** | ** | **-0.16** | *** | **-0.15** | ** | **-0.07** | |
| PC6 | -0.01 | | -0.05 | | -0.04 | | -0.09 | | -0.07 | | -0.08 | |
| PC8 | 0.04 | | **-0.14** | ** | -0.07 | | -0.06 | | -0.08 | | -0.03 | |
| PC9 | -0.03 | | -0.05 | | -0.04 | | -0.02 | | -0.01 | | -0.07 | |
| PC10 | **-0.11** | * | 0.05 | | -0.03 | | 0.00 | | -0.06 | | 0.06 | |
| PC12 | **0.11** | * | -0.07 | | -0.01 | | 0.01 | | -0.02 | | -0.09 | |
| PC15 | 0.05 | | 0.00 | | -0.01 | | 0.00 | | **0.11** | * | -0.03 | |
| PC19 | **0.28** | *** | 0.06 | | **0.10** | * | 0.10 | * | 0.09 | | -0.06 | |
| PC26 | 0.01 | | 0.08 | | 0.01 | | 0.00 | | 0.02 | | 0.00 | |
| PC29 | 0.00 | | 0.07 | | 0.04 | | 0.02 | | 0.02 | | 0.09 | |
| PC32 | **-0.18** | *** | -0.02 | | **-0.12** | * | -0.08 | | **-0.14** | ** | 0.03 | |
| **PC33** | **-0.23** | *** | **-0.20** | *** | **-0.17** | *** | **-0.17** | *** | **-0.15** | ** | 0.00 | |
| PC41 | **0.09** | * | -0.08 | | 0.01 | | -0.03 | | 0.02 | | -0.07 | |
| PC391 | 0.03 | | -0.03 | | 0.05 | | 0.02 | | 0.04 | | 0.03 | |
| **Overall** | **0.59** | *** | **0.15** | *** | **0.28** | *** | **0.27** | *** | **0.32** | *** | 0.03 | |
| **Acceleration** | -0.03 | | **0.15** | *** | **0.13** | ** | **0.17** | *** | **0.16** | *** | **0.12** | * |

| BH Corrected P-values | |
|---|---|
| > 0.05 (n.s.) | |
| < 0.05 | * |
| < 0.005 | ** |
| < 0.0005 | *** |

#### Table S3. Linear Models for PCBrainAge.C Acceleration

| | Testing | | PFC | | ST | | CBM | |
|---|---|---|---|---|---|---|---|---|
| **predictor** | **Estimates** | **p** | **Estimates** | **p** | **Estimates** | **p** | **Estimates** | **p** |
| **(Intercept)** | 33.33 | <0.001 | 54.76 | <0.001 | 48.05 | <0.001 | 47.20 | <0.001 |
| **Age** | 0.46 | <0.001 | 0.39 | <0.001 | 0.46 | <0.001 | 0.17 | <0.001 |
| **Prop N** | 3.88 | 0.138 | -18.58 | <0.001 | -16.80 | <0.001 | -18.94 | 0.031 |
| **Observations** | 700 | | 333 | | 333 | | 333 | |
| **R2/R2 adjusted** | 0.389/0.388 | | 0.254/0.250 | | 0.310/0.306 | | 0.085/0.080 | |

#### Table S4. Linear Mixed Effect Models in Multi-Region Brain Data Improves upon OLS Regression

| | OLS Regression | | LME Model | |
|---|---|---|---|---|
| **Predictors** | Estimates | p | Estimates | p |
| (Intercept) | 63.79 | <0.001 | 49.03 | <0.001 |
| Age | 0.35 | <0.001 | 0.34 | <0.001 |
| Prop N | -35.68 | 0.138 | -15.96 | <0.001 |
| **Random Effects** | | | | |
| $\sigma^2$ | | | 18.48 | |
| $\tau_{00}$ | | | $169.63_{region}$ | |
| ICC | | | 0.90 | |
| N | | | $3_{region}$ | |
| **Observations** | 997 | | 997 | |
| **$R^2$/$R^2$ adjusted** | 0.325/0.324 | | 0.030/0.905 | |

### Supplemental Figures

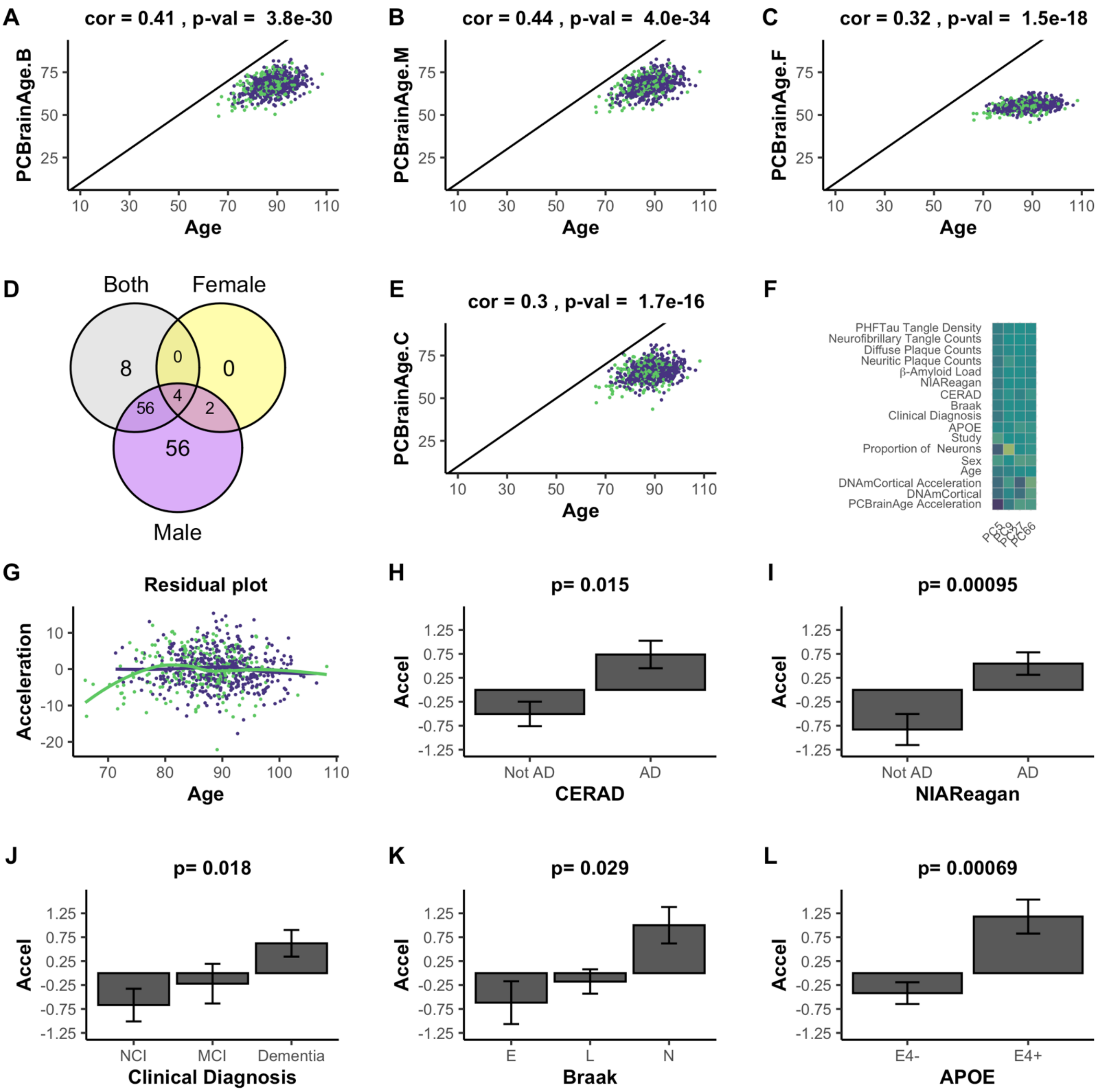


**Figure S1: Removing Schizophrenia Patients from Training Sample Doesn't Improve Model Performance**. When sfchizophrenia patients are not included in the training data for each model, performance is significantly reduced in all iterations (**A-C**) when compared to the original training models (**Figure 1D-F**). Core PCs consist of just 4 elements (**D**) which do not predict age as well in the retrained core model (**E**) as in the original (**Figure 1G**). The PCs selected are quite similar to those of the original model (**F**), but do not capture components of AD in the test dataset with age acceleration as strongly as in the original PCBrainAge Core model (**H-L**).

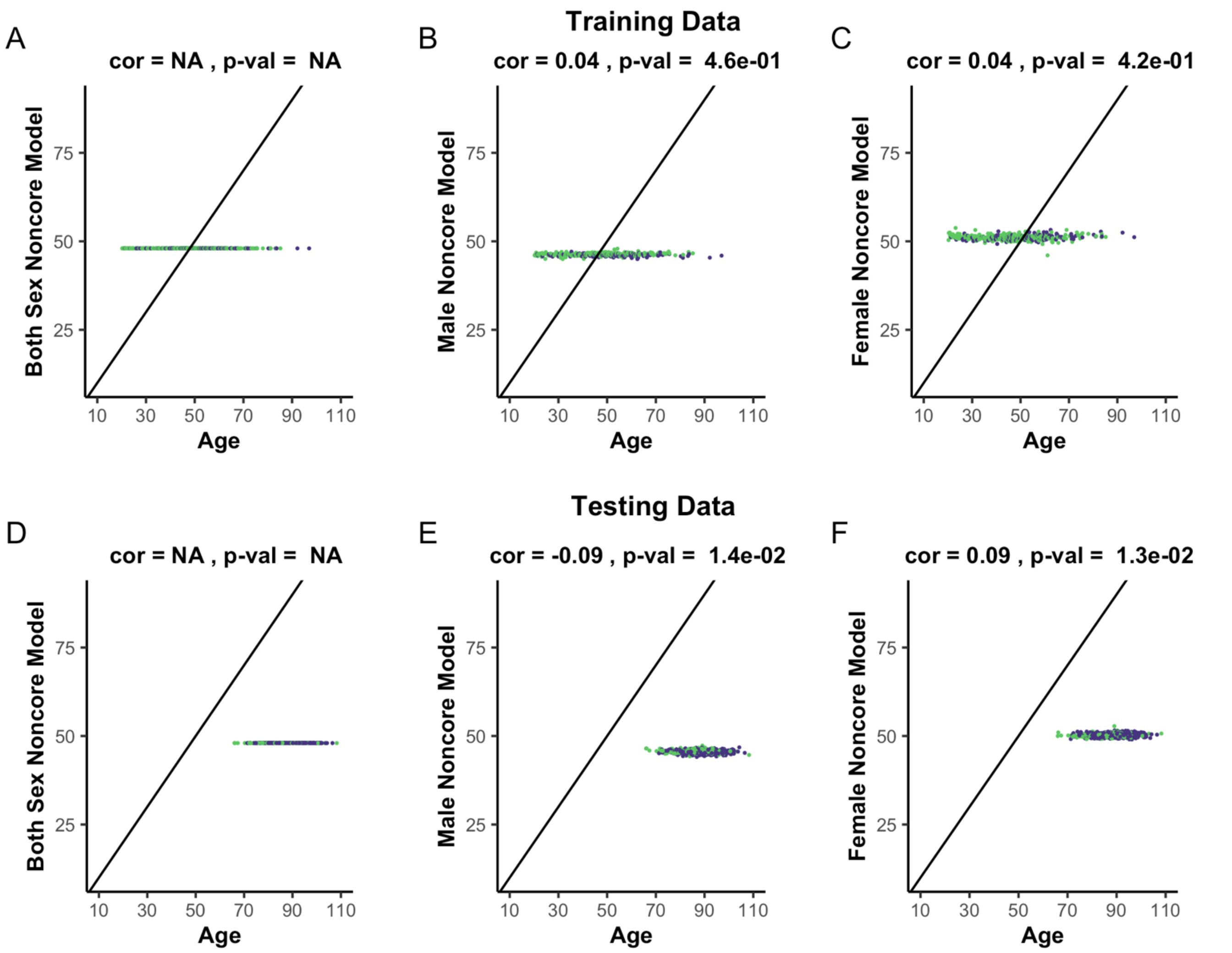


**Figure S2: Confirmation that Core PCs Contain the Aging Signal.** All panels from this figure are comparable to those of Figure 1. The major difference in the present figure is that the principal components in the core of Figure 1G have been removed from consideration for the elastic net model training. Clearly, the models are unsuccessful at predicting age.

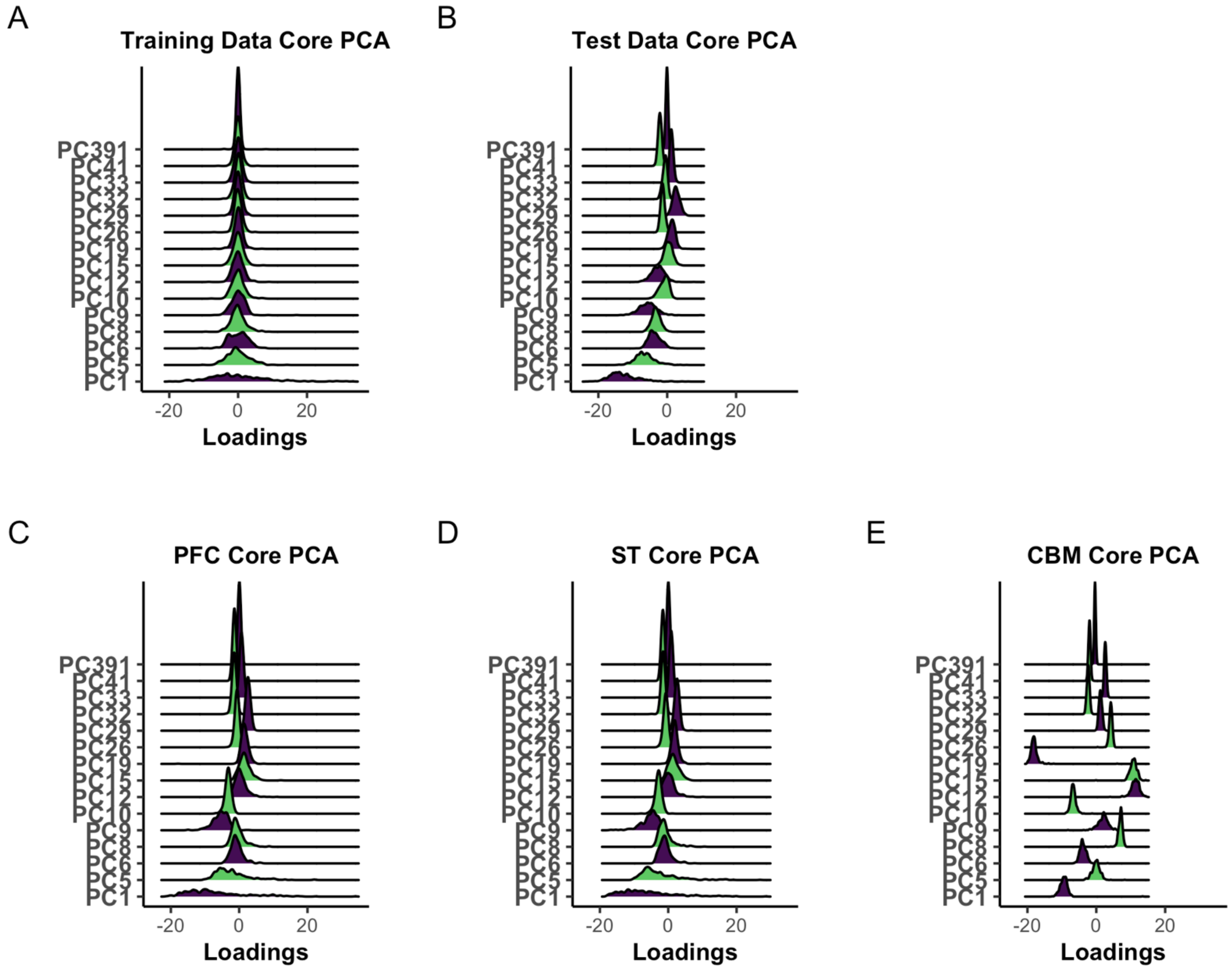


**Figure S3: All dataset sample loading distributions.** The distribution of loadings for each core PC is evaluated in the training dataset, across all ages (**A**) as compared to Figure 2C, in the testing dataset (**B**) exactly as Figure 2D, and in the multi-region dataset in PFC (**C**), ST (**D**), and CBM (**E**).

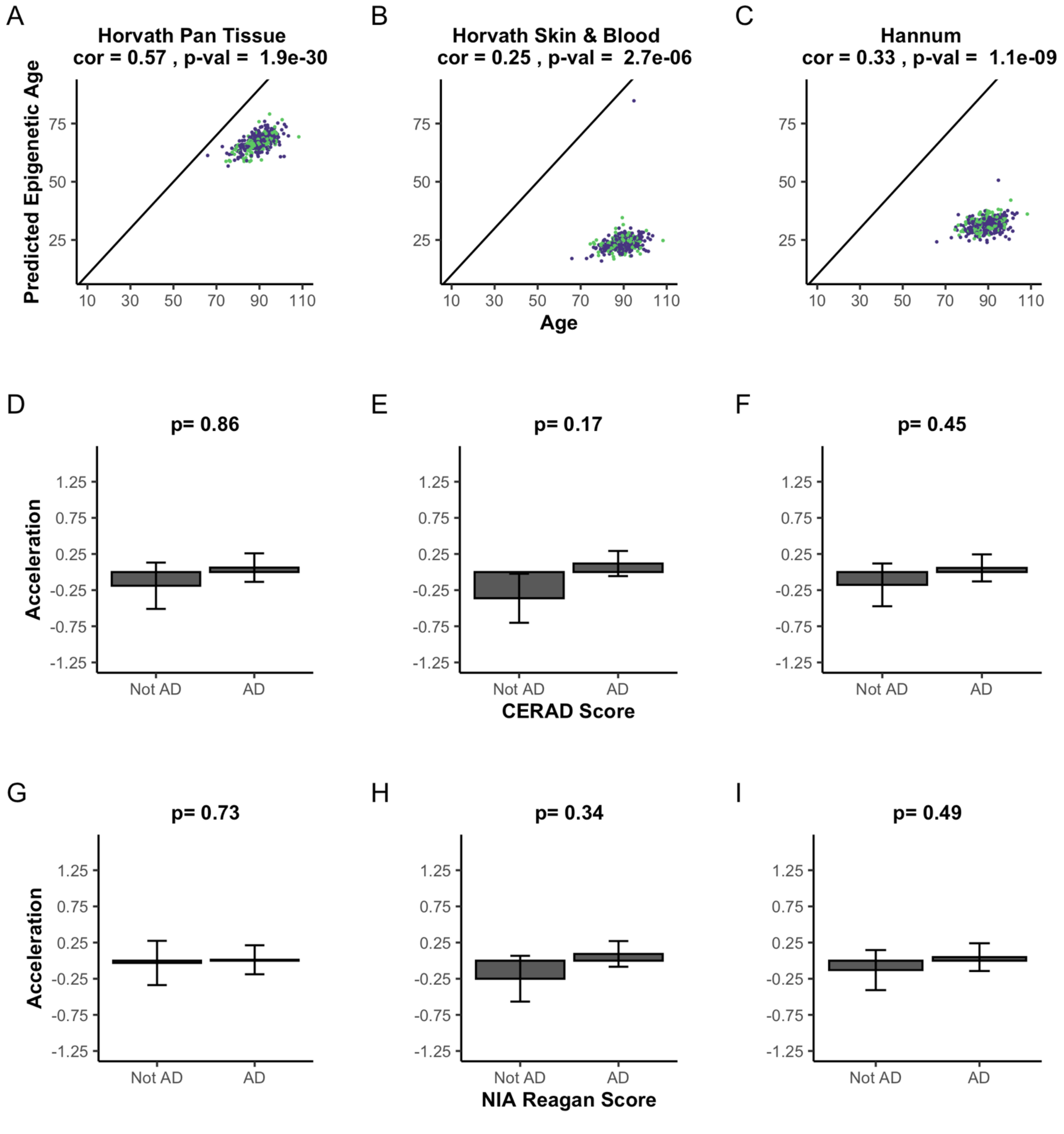


**Figure S4: Existing Epigenetic Clocks do not Robustly Capture Hallmarks of AD in Cerebellum.** The 3 most used epigenetic clocks, the Horvath pan tissue (**A**, **D**, **G**), Horvath Skin & Blood (**B**, **E**, **H**), and the Hannum clock (**C**, **F**, **I**) were calculated in our cerebellum dataset. The predicted ages have low correlation with age at death (**A-C**). Age acceleration, calculated as the residuals of a linear model of the clocks' predictions onto age and proportion of neurons were not significantly correlated with CERAD scoring criterion (**D-F**). Age acceleration for all three clocks were also not correlated with combined criterion NIA Reagan scores (**G-I**).

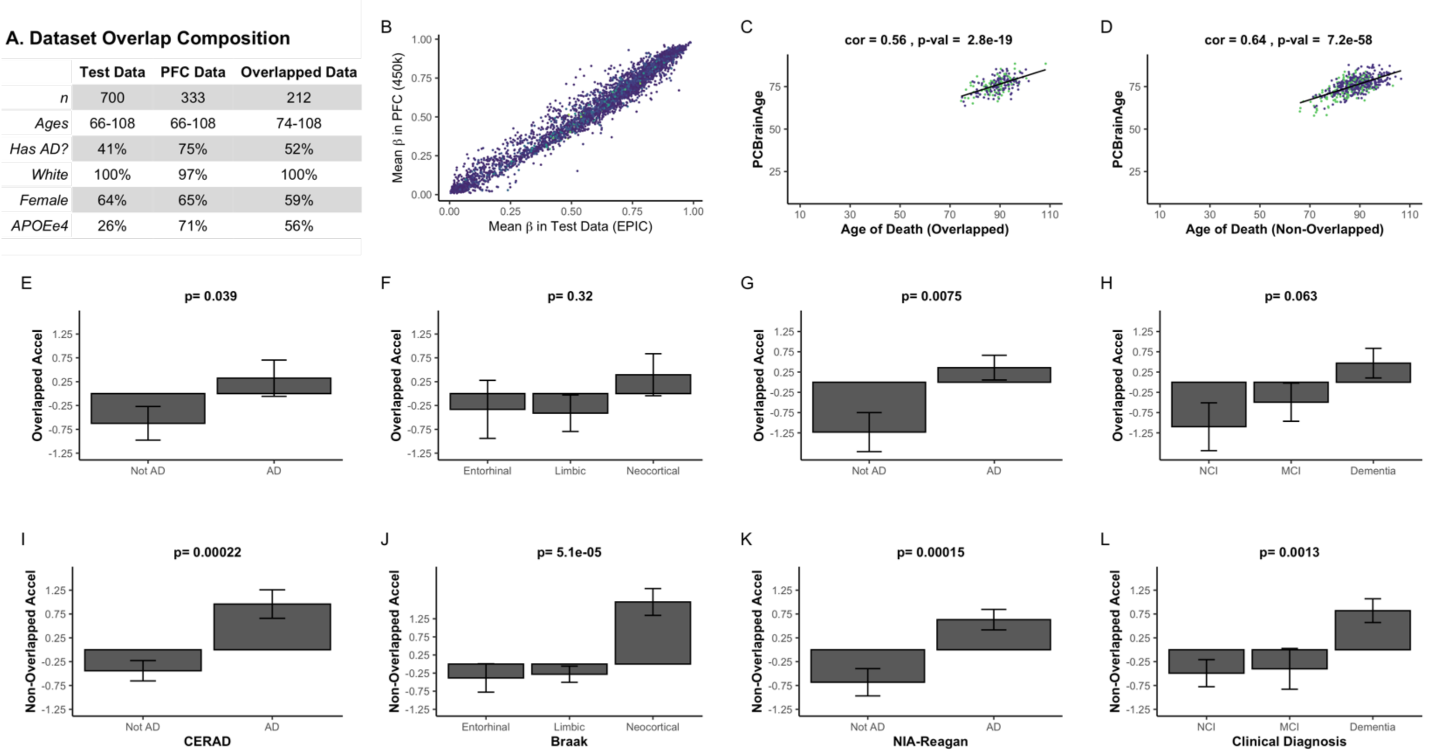

A. Dataset Overlap Composition

| | Test Data | PFC Data | Overlapped Data |
|---|---|---|---|
| *n* | 700 | 333 | 212 |
| *Ages* | 66-108 | 66-108 | 74-108 |
| *Has AD?* | 41% | 75% | 52% |
| *White* | 100% | 97% | 100% |
| *Female* | 64% | 65% | 59% |
| *APOEe4* | 26% | 71% | 56% |



**Figure S5. Differences in Overlapped Versus Non-Overlapped Individuals From Two Datasets.** The composition of the 2 datasets and the subset of overlapped individuals is described (**A**). The CpGs which are used to generate the principal components' weights were generated by multiplying PC model weight by the CpG loadings across the PCs, and summing the total weight for each CpG. The weights were converted to z-scores, and those with a value of 3 or higher (4582 CpGs) are plotted to compare the agreement of mean beta values in both datasets (**B**). When weights of z-score < 3 are plotted, points occupy a similar space but depart further from the diagonal. Overlapped individuals (**C**) show similar correlations of PCBrainAge prediction to age as the non-overlapped group (**D**). The two groups' PCBrainAge acceleration was further correlated to CERAD scores (**E**, **I**), Braak scores (**F**, **J**), NIA Reagan scores (**G**, **K**), and cognitive diagnoses (**H**, **L**). While separation is clearer when using non-overlapped individuals (**I**, **J**, **K**, **L**), this may be due to the larger set of individuals.
